# Molecular Cloning and Biochemical Characterisation of a Novel Acidic Laminarinase Derived from Jermuk Hot Spring Metagenome

**DOI:** 10.1101/2024.01.22.576621

**Authors:** Ani Paloyan, Mariam Karapetyan, Hasmik Grigoryan, Anna Krüger, Christin Burkhardt, Garabed Antranikian

## Abstract

Laminarinase, an enzyme with a specific affinity for laminarin—a complex polysaccharide found in the cell walls of brown algae and select marine organisms—was investigated in this study. We cloned and characterised a gene encoding a putative glycoside hydrolase family 16 (GH16) laminarinase from the Jermuk hot spring metagenome by heterologous expression in *Escherichia coli*. The resulting product, named Jermuk-LamM, represents a novel endo-1,3-β-D-glucanase (EC 3.2.1.39) with only 48.1 % amino acid sequence similarity to previously characterised GH16 family members catalogued in the NCBI database. To date, this stands as the sole described endo-1,3-β-D-glucanase within the *Marinimicrobia* phylum.

Jermuk-LamM, identified as an acidic laminarinase, exhibits robust enzymatic activity at pH 5.0 and a temperature of 55 °C, maintaining its function for a duration of at least 7 hours. Notably, this enzyme effectively catalyses the hydrolysis of both soluble and insoluble (1,3)-β-D-glucans, as well as (1,3;1,4)-β-D-glucans, displaying a pronounced preference for laminarin. The specificity of Jermuk-LamM lies in its cleavage of 1,3-β-D-glucosidic linkages, yielding monosaccharides, disaccharides, and oligosaccharides. These breakdown products hold the potential for conversion into energy carriers, including alcohols, methane, and hydrogen.

The enzyme’s exceptional specific activities, coupled with its resistance to various additives, render Jermuk-LamM a promising candidate for various industrial applications, encompassing the realms of biofuel and pharmaceutical production.

## Introduction

Polysaccharides with β-1,3-linkages, such as laminarin, curdlan, lichenin, β-glucan, have been found in animals, plants, algae and microorganisms. They have diverse functional properties and play an essential role in sustaining life. Laminarin, consists of glucose units with β-1,3-linkages and partial branches connected through β-1,6-linkages [1]. It serves as the primary storage polysaccharide in macroalgae [2]. Furthermore, it is also a component of the cell walls in most fungal species [3]. Curdlan is unbranched and consists of β-1,3-linked glycosyl residues with a chain length of about 135 residues [4, 5], sometimes with a few intra- or interchain 1,6-linkages [6]. It is an extracellular and capsular polysaccharide produced by a variety of soil-dwelling bacteria [7, 8]. Lichenin, a major polysaccharide component of *Cetraria islandica* (Iceland moss) is composed of β-D-glucopyranose units linked in a linear manner by β-1,4 and β-1,3 glycosidic bonds [9, 10] and plays a central role in forming symbiotic relationships with algae in lichens. The other known source of polysaccharides of this type is β-glucan, a major component of endosperm cell walls and sub-aleurone layer of Gramineae, consisting of glucose linked by both β-1,4 and β-1,3 glycosidic linkages [11, 12].

Unlike starch, these polysaccharides are non-food-based sugar sources and potentially more suitable resources for biofuels, as well as for pharmaceuticals, cosmetics packaging material, etc [13, 14]. Recently, increasing attention is being drawn to macroalgae as an ideal biomass source with high levels of carbohydrates with low to zero lignin content [15]. Moreover, macroalgae do not compete with agricultural land use [16]. Therefore, it is important to design biocatalysts to develop a cost-competitive process for macroalgae carbohydrates utilization [17, 18, 19]. In macroalgae laminarin content ranges from 1-25% of total weight located in vacuoles [20].

Laminarinase, also known as laminarin hydrolase or β-1,3-glucanohydrolase catalyses the hydrolysis of β-1,3-glucosidic bonds except the LamR from *Rhodothermus marinus* [21] and SCLam derived from the soil metagenome [22], which are able to cleave as 1,3-as well as 1,4-β-D-glycosyl linkages. Laminarinases have garnered interest due to their potential applications in various industries, including biofuel production, food processing, and pharmaceuticals [23, 24, 25, 26].

Laminarinases are classified into exo-β-1,3-glucanases (EC 3.2.1.58) and endo-β-1,3-glucanases (EC 3.2.1.39 and EC 3.2.1.6) [27]. Eendo-β-1,3-glucanases hydrolyze β-1,3-glucans and release glucose residues or oligosaccharides whereas exo-β-1,3-glucanases cleave glucose residues from the non-reducing end of the substrate. Laminarinases have been characterised from plants [28], fungi [29, 30, 31, 32], algae [33], *Actinobacteria* [34, 35], Archaea [36], soil bacteria [37, 38, 39], thermophilic [40, 36, 41, 42, 43, 44] and marine bacteria [45, 46, 47, 21, 48, 49]. Most of the characterised laminarinases belong to glycosyl hydrolase family 16 (GHF16) [50].

Considerable attention has been devoted to the characterisation and continuous improvement of laminarinases, aiming to establish a cost-competitive approach for converting laminarin into fermentable sugars [51, 52].

Research findings have confirmed that metagenomics is a promising approach to tap into the genetic diversity of microbial communities and to discover novel enzymes including several novel glycoside hydrolases belonging to different GH families [53, 54, 55, 22]. A metagenome obtained from Jermuk hot spring served as a source for laminarinase identification. Jermuk hot springs are located within the geothermal system of the Lesser Caucasus mountains at about 2080 meter altitude above sea level, the temperature of water of the hot springs water varies from 40 to 53 °C. Different strains belonging to *Bacillus, Parageobacillus, Geobacillus* and *Anoxybacillus* genera have already been isolated from Jermuk hot springs, producing valuable sources of hydrolases [56]. This study offers an in-depth characterisation of a laminarinase derived from a metagenome of Jermuk hot springs. This work describes the first characterised enzyme from the *Marinmicrobia* phylum. The potential application of this enzyme for bioconversion of natural polysaccharides was evaluated.

## Materials and methods

### Reagents and substrates

Phusion® DNA Polymerase, BsaI restriction enzyme and T4 DNA ligase, were purchased from New England Biolabs (Hitchin, UK). Isopropyl β-D-1-thiogalactopyranoside (IPTG) was purchased from Merck, UK. The molecular weight marker for SDS PAGE was purchased from Thermo Fisher Scientific (Cramlington, UK). Carboxymethyl cellulose (CMC) and lichenin were obtained from Merck and β-glucan (barley), curdlan (*Alcaligenes faecalis*), as well as AZCL substrates from Megazyme. Standards, laminaritetraose, laminaritriose, laminaribiose were purchased from from Sigma, glucose from Merck.

### Bacterial strains, plasmids, metagenome

The metagenome DNA obtained from Jermuk hot spring was used for ORFs amplification. Samples were collected from different locations of a Jermuk hot spring coordinates (39.8411693, 45.6678023), temperature 40-45 °C, pH 8-9 (sample type: water and sediment). The collected material was transported and stored at 4 °C.

The DNA was isolated using the PowerSoil Isolation Kit by MO BIO Laboratories Inc. according to the manufacturer’s instructions. The obtained DNA was then subjected to Illumina MiSeq sequencing. Sequence data processing, assembly and annotation were performed as previously described [57].

pET28 plasmid modified for Golden Gate cloning [58], *E. coli* TOP10 and *E. coli* BL21 CD3 (DE3) strains were used for gene cloning and expression.

### Sequence analysis, cloning of ORFs

A DLam gene from *Dictyoglomus thermophilum* was used to scan contigs for predicted ORFs. Potential ORFs encoding putative endo-β-1,3-D-glucanases were identified and reannotated contigs were compared to the NCBI (Blastx) database. To identify the most related nucleotide and amino acid sequences for ORFs, BLASTn, BLASTp, and PSI-BLAST analysis were performed. The prediction of functional and structural domains, catalytic sites and signal sequences were performed with applications such as Conserved Domain Database (CDD) [59], ScanProsite [60], and SignalP 4.0 [61], SMART [62]. For multiple sequence alignment and construction of a phylogenetic tree from the GenBank database, amino acid sequences of characterised β-1,3-glucanases and β-1,3(4)-glucanases of the family GH 16 were selected. Sequence alignment was prepared using Multalin [63] and figures prepared with ESPript [64]. The phylogenetic tree was constructed by using MEGA11, calculated by the neighbor joining method [65]. The evolutionary history of the Jermuk-LamM was inferred by using the Maximum Likelihood method and JTT matrix-based model [66]. The highest log likelihood is −17122.20. Initial tree(s) for the heuristic search were obtained automatically by applying Neighbor-Join and BioNJ algorithms to a matrix of pairwise distances estimated using the JTT model, and then selecting the topology with superior log likelihood value. This analysis involved 16 amino acid sequences.

Structure of Jermuk-LamM was generated using the Phyre2 server [67] visualized by Mol*3D viewer [68]. The theoretical molecular weight (MW) and isoelectric point (pI) were estimated using the ExPASy, ProtParam tool Compute MW/pI [69].

Benchling was used for specific primer design, *in silico* PCR and assembly experiments. For cloning, Jermuk-lamM gene was amplified using LamM-F (5’**GACGGTCTCTAATG**CCCGAAGACGAATCGCCTCAGG3’) and LamM-R (5’**GTCGGTCTCTACCT**GGATTTGATCCGCTGGAAGATACGGAC3’) primers (primer extending sequences are indicated in bold) within the following PCR conditions: 98 °C for 1 min followed by 30 cycles of 98 °C for 30 s, 65 °C for 30 s and 72 °C for 1 min, followed by a final elongation at 72 °C for 10 min, using the metagenomic DNA as template. After examination by electrophoresis the amplified product was purified with the QIAquick PCR Purification Kit, then assembled via one-pot Golden Gate cloning [58] and CIDAR MoClo system [70], and used to transform *E. coli* TOP10 cells. Recombinant plasmid DNA was extracted from insert-positive clones and after sequence confirmation used to transform *E. coli* C43(DE3) or *E. coli* BL21(DE3).

### Expression of the Jermuk-lamM gene and enzyme purification

For protein expression *E. coli* C43(DE3) harboring the plasmids pET28GGLacZ::*Jermuk-lamM* was grown in LB media (35 μg/mL kanamycin) at 37 °C and 160 rpm. Gene expression was induced at OD_600_ =0.5-0.6 by adding IPTG to a final concentration of 1 mM. Cells were harvested by centrifugation at 9000 × *g* at 4 °C for 20 min. The resulting cell pellet was stored at −20 °C.

For purification, 0.2 g cells were resuspended per 1 mL lysis buffer (50 mM NaH_2_PO_4_, 300 mM NaCl, 10 mM imidazole, pH 8) and disrupted by three passages through a French pressure cell with constant pressure of 1,000 psi (French Pressure Cell Press, SLM-Aminco). Cell debris was removed by centrifugation (20,000 × *g*, 4 °C, 30 min) and supernatant was loaded onto a 1 ml Ni-NTA Superflow column (Qiagen). Proteins were eluted by an increasing imidazole gradient according to the manufacturer’s instructions. Eluted fractions were pooled, washed three times with buffer A (50 mM Na-phosphate buffer, pH 7.2, 150 mM NaCl) by ultrafiltration in an Amicon filter unit (Amicon Ultra-15, 1000 MWCO, Merck Millipore). For final purification via size exclusion chromatography, protein solutions were loaded onto a Sephacryl S-100 column previously equilibrated with buffer A. Protein fractions containing the purified enzymes were pooled and stored at 4°C.

Protein samples were analyzed by SDS-PAGE (12.5 %) [71]. Protein concentration was determined according to Bradford [72], with bovine serum albumin as standard.

### Characterisation of Jermuk-LamM

Unless otherwise noted the standard assay was carried out at 65 °C for 10 min in 500 μL reaction mixture using 0.25 % (*w/v*) laminarin from *Laminaria digitata* (Merck) as substrate in 50 mM sodium phosphate buffer pH 6.0 (adjusted at assay temperature), and 5.0±0.5 μg (0.11 mg/mL) enzyme sample.

Additionally, blanks without enzyme were performed by default for all measurement series. The hydrolytic activities of the purified enzymes were detected by measuring the reducing sugars with 3,5-dinitrosalicylic acid (DNS) according to Miller [73] with glucose as standard. In brief, after enzyme reaction 500 μL reaction mixture were mixed with 500 μL DNS reagent (1 % (*w/v*) DNS, 30 % (*w/v*) potassium sodium tartrate, 0.4 M NaOH) and were incubated for 5 min at 100 °C. Samples were subsequently cooled on ice to room temperature and absorption was measured at 546 nm. All measurements were done in triplicates. One unit of enzyme activity was defined as the amount of enzyme required to release 1 μmol of reducing sugars per minute.

The influence of temperature on enzyme activity was examined by performing the standard assay at temperatures from 30 to 80 °C. To investigate the temperature sensitivity, the enzyme samples were pre-incubated with a concentration of 0.1 mg/mL in the same range of temperature for 10 minutes. Subsequently, they were stored on ice and residual activities were measured by using the standard assay. To investigate the temperature stability at selected temperatures Jermuk-LamM was pre-incubated at 55, 60, and 65 °C, respectively, and samples were taken in time intervals up to 24 h. Residual activities were measured by using the standard assay. To determine the influence of the pH on enzyme activity, a standard assay was performed using Britton-Robinson buffer (50 mM) in a range of pH 2-10 in the reaction mixture [74].

The influences of metal ions on enzyme activity were analyzed by using a standard assay after incubating the enzyme in the presence of 5 mM AlCl_3_, CaCl_2_, CoCl_2_, CrCl_3_, CuCl_2_, FeCl_2_, FeCl^3^, KCl, MgCl_2_, NaCl, NiCl_2_, RbCl, SrCl_2_ or ZnCl_2_ for one-hour at room temperature. The influences of additives such as 3-((3-cholamidopropyl)dimethylammonio)-1-propanesulfonate (CHAPS), SDS, Triton X-100, Tween 20, Tween 80, guanidine hydrochloride, urea, dithiothreitol (DTT), β-mercaptoethanol, EDTA, iodoacetic acid, Pefabloc, cetyltrimethylammonium bromide (CTAB) and sodium azide were examined by the standard assay procedure followed by one-hour incubation at room temperature. All additives were tested at a concentration of 5 mM under standard conditions. A blank experiment was performed for each metal and additive.

For analysing the substrate spectrum, the specific activities of Jermuk-LamM were measured using different substrates. All substrates were used at a final concentration of 0.25 % (*w/v*). In case of curdlan an undissolved and a dissolved (amorphous) form were tested. To achieve an amorphous type of curdlan, 0.2 g of the substrate were first solubilized in 6 mL alkaline solution (0.6 M NaOH) and subsequently neutralized with HCl to a concentration of 0.5 % (*w/v*) and pH 6.0 in 50 mM sodium phosphate buffer. A blank experiment was conducted for each substrate.

### Determination of the Hydrolysis Products

For determination of the hydrolysis products 0.25 % (*w/v*) substrate were incubated with Jermuk-LamM in standard reaction mixtures at 60 °C for 18 h. After the inactivation of the enzyme samples at 100 °C for 10 min, samples were centrifuged (20.000 × *g*, 10 min) and the supernatant was filtered using a 0.22 μm membrane filter unit. Hydrolysis products were analyzed by high-performance liquid chromatography (HPLC) under the following conditions: Aminex HPx-42A column (Bio Rad), 80 °C, 0.6 mL/min flow rate, water as mobile phase, HPLC 1260 Infinity II LC System (Agilent Technologies) with RI detector. Laminaritetraose, laminaritriose, laminaribiose, and glucose were used as standards.

### Nucleotide sequence accession number

The sequence of *Jermuk-lamM* was deposited in GenBank (accession number OK490392).

## Results

A total of 68 ORFs was identified in the Jermuk metagenome dataset. After cloning and initial plate tests, recombinant *Jermuk-lamM* was selected for further analysis. The deduced enzyme showed activity on agarose plates using AZCL-Curdlan, AZCL-Barley β-glucan and AZCL-Pachyman as substrate (data not shown).

The ORF *Jermuk-lamM* was identified by a sequence-based screening of a metagenome obtained from a hot spring of Jermuk, Armenia. The ORF was found to encode a putative glycoside hydrolase family 16 enzyme. It consisted of 783 bp, with the encoded endo-β-1,3-glucanase consisting of 287 amino acids. The premature Jermuk-LamM was evaluated to be an extracellular protein with the signal peptide sequence MPKLYFYLAGALLALNLLMTCHEPSKA at the N terminus that is cleaved between Ala27 and Pro28, as predicted by the SignalP 5.0 server. Domain prediction revealed that Jermuk-LamM consists of one single catalytic domain GH16 without any further known structural elements except a signal peptide which is connected to the catalytic domain by a long amino acid linker sequence (PEDESPQDTIRGPVNYQLVWHDEFNDSTIDLNKWIFEVNAH) (**Figure S1**). A theoretical molecular mass of 34.9 kDa and an isoelectric point (pI) of 5.47 were calculated for the premature enzyme. Both the nucleotide and protein sequence of recombinant Jermuk-LamM had less than 75 % identity with all glycoside hydrolases deposited in the NCBI database. According to the BLAST data the encoded protein (Jermuk-LamM) showed 71.3 % (query coverage 100 %) identity to GH16 family protein of *Candidatus Marinimicrobia* bacterium (GenBank: HDP67611.1), 67.2 % (query coverage 100 %) identity to GH16 family protein candidate division KSB1 bacterium (GenBank MBD3289618.1), 65.84 % (query coverage 98 %) identity to GH16 family protein of *Calditrichaeota bacterium* (GenBank: RMF58681.1), all of which are uncultivable marine bacteria and none of the mentioned genes has been cloned or their encoding proteins have been characterised so far. Phylogenetic analysis indicated that Jermuk-LamM forms a cluster with endo-1,3(4)-beta-glucanase originating from *Acetivibrio thermocellus* and is more distantly related to endo-beta-1,3-glucanase from the hyperthermophilic Archaeon *Pyrococcus furiosus* DSM 3638 and laminarinase of the extreme thermophilic bacterium *Thermotoga maritima* MSB8 (**Figure 1**). Interestingly, in contrast to Jermuk-LamM endo-1,3(4)-beta-glucanase (Lic16A) of *Acetivibrio thermocellus* (previously known as *Clostridium thermocellum*) is a complex protein consisting of a number of structural modules including a catalytic domain GH16 [75]. Moreover, in terms of the overall amino acid sequence, Lic16A (CAC27412.2) exhibits a relatively low identity (11 %) to Jermuk-LamM while laminarinase of *Thermotoga maritima* MSB8 (AAD35118.1), being in a more distant cluster, has the highest (48.1 %) amino acid sequence identity to Jermuk-LamM․ However, when focusing solely on the identity of the CH16 catalytic domains, the scenario shifts. The catalytic domains of Lic16A and Jermuk-LamM share the highest identity with 56 %.

**Figure 1.**
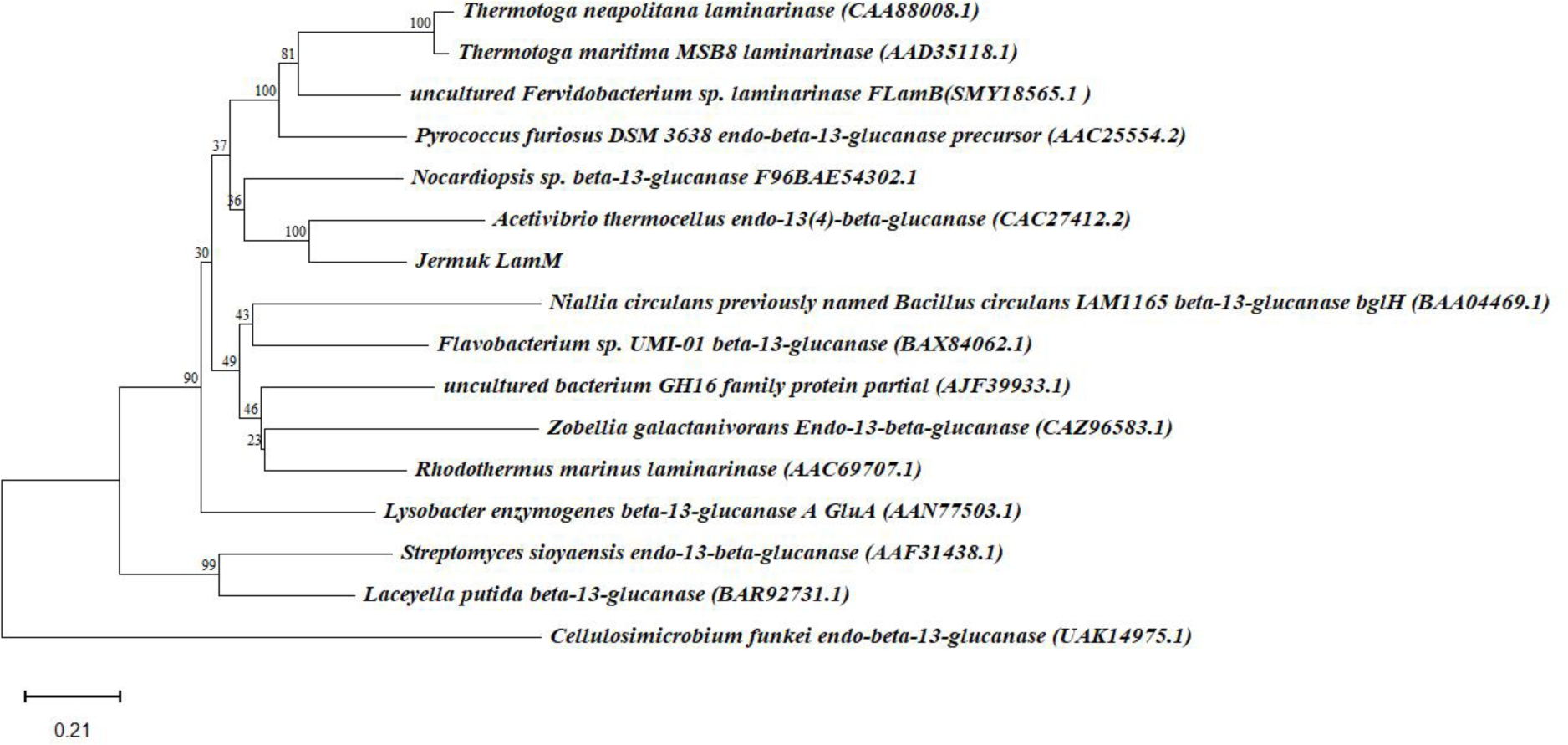
Phylogenetic tree of Jermuk-LamM enzymes. The evolutionary history was inferred by using the Maximum Likelihood method and JTT matrix-based model. The tree is drawn to a scale, with branch lengths measured in the number of substitutions per site (next to the branches). The tree with the highest log likelihood (−17122.20) is shown.

Structure predictions for Jermuk-LamM revealed a sandwich-like β-jelly roll topology, which is typical for laminarinases of class GH16. It comprises 18 β-strands, and an α-helix (**Figure S2**). The predicted structure of Jermuk-LamM was aligned with those of the catalytic domain of laminarinases from *T. maritima* (TmLamCD), alkaliphilic *Nocardiopsis* sp. strain F96 (BglF), *Rhodothermus marinus* (RmLamR), and *Pyrococcus furiosus* (pfLamA), all of which have been structurally characterised (Figure 2). Protein sequence alignment analysis of Jermuk-LamM using the Blast program shows an identity of 52 % with BglF (PDB code 2HYK) [34], 48 % with TmLamCD (PDB code 3AZY) [76], 45 % with pfLamA (PDB code 2VY0) [77] and 44 % with RmLamR (PDB code 3ILN).

**Figure 2.**
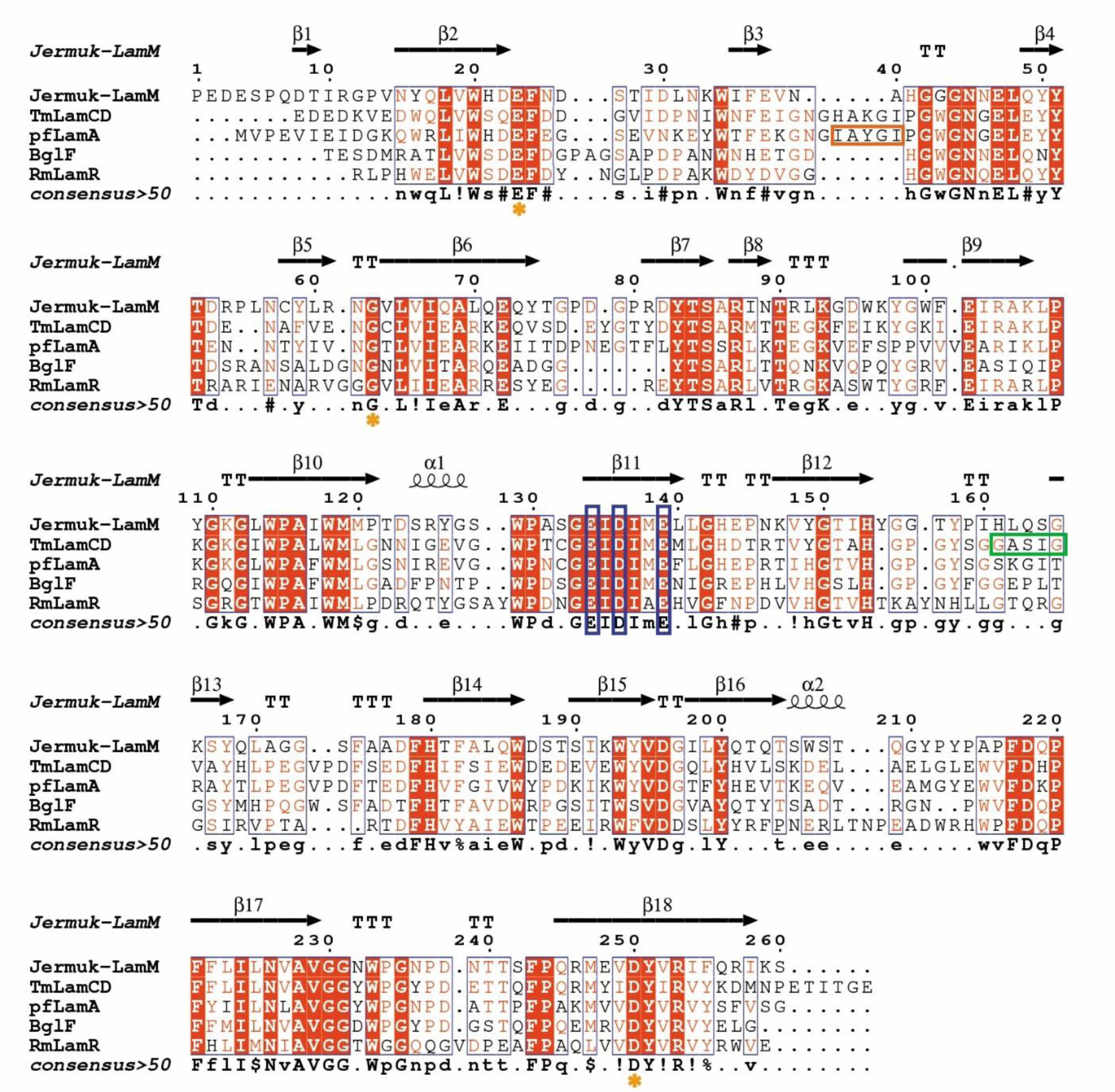
Annotated sequence alignment of Jermuk-LamM enzymes. Multiple sequence alignment of catalytic domain of Jermuk-LamM, BglF (PDB:2hyk), TmLamCD (PDB: 3azy), pfLamA (PDB: 2vy0) and RmLamR (PDB: 3iln) was performed using Multalin and visualised with ESPript. Secondary structure elements from the predicted crystal structure of the Jermuk-LamM are shown above the alignment. Strictly conserved residues are shown with a red background, while partially conserved residues are shown with red text. Conserved residues involved in metal coordination are highlighted with orange asterisk. Conserved catalytic (Glu, Asp, Glu) triad is highlighted in blue box. “GASIG” flexible loop of TmLamCD is highlighted in green box. “Kink” peptide of pfLamA is highlighted in orange box. The sequence identity between Jermuk-LamM and BglF, between Jermuk-LamM and TmLamCD, Jermuk-LamM and pfLamA and Jermuk-LamM and RmLamR are 52 %, 48 %, 45 % and 44 %, respectively.

Despite the high identity between the catalytic domains of Jermuk-LamM and TmLamCD, certain differences were observed in terms of specific structural elements that hold functional significance. Notably, one of these elements has been described a “GASIG” flexible loop in TmLamCD which is lacking in the predicted structure of Jermuk-LamM. It has been proven that this loop together with Trp232 has a significant role in the regulation of endo- or exo-activity of the enzyme and a preference to release laminaritrioses in long-chain carbohydrate hydrolysis. Meanwhile, Arg-85 which has been shown to have a critical role in β-1,3-glucan substrate selection was found to be conservative for Jermuk-LamM [76]. Regarding pfLamA, its structure revealed a kink of six residues (72–77) at the entrance of the catalytic cleft, which was shown to be important for substrate preference and ensured higher activity to laminarin than lichenin. This particular peptide is notably missing in Jermuk-LamM, as well as in BglF and RmLamR. Interestingly, a similar peptide is present in TmLamCD, although its specific functional role remains undocumented. The sequence EIDIME, which includes the catalytic glutamate residues of the active site conserved within the GH16 family, was identified between residues Gly160 and Leu167 for the Jermuk-LamM protein.

### Overexpression of the gene *Jermuk-lamM* in *E. coli* and purification of the recombinant enzyme

The ORF Jermuk-lamM without signal peptide was amplified with attached restriction recognition sites omitting start and stop codon and ligated into the modified vector pET28, which includes a 6xHis tag encoding region. The expression of the Jermuk-LamM construct in *E. coli* BL21 (DE3) using common expression conditions (such as induction by 1 mM IPTG for 4 hours at 37 °C) resulted in the formation of inclusion bodies. Thus, more than 80 % of the protein remained in the pellet. To overcome this problem a number of parameters such as the concentration of IPTG, induction time and temperature, as well as different expression hosts were tested. The expression conditions were optimized, resulting in optimal protein production when expressed in *E. coli* C43(DE3) at 30 °C for 24 hours (data not shown). The obtained protein harboring a C-terminal 6xHis affinity tag was purified via affinity chromatography and subsequent size exclusion chromatography. Jermuk-LamM was purified to homogeneity with a 228.73 U/mg specific activity and a final yield of 13.6 % (**Table S1**). The SDS-PAGE revealed a molecular weight of approximately 30 kDa, which was in accordance with the predicted molecular size of 29.8 kDa (without signal peptide) (Figure 3).

**Figure 3.**
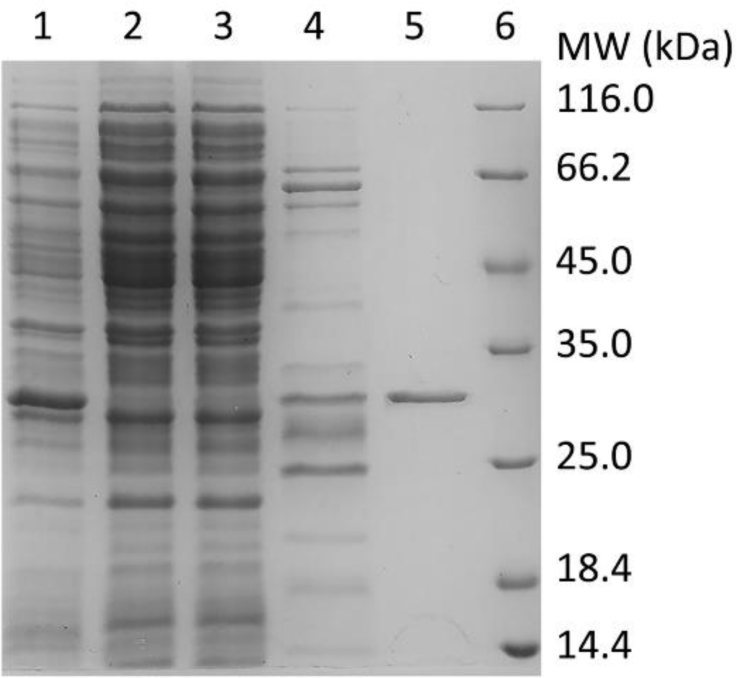
SDS-PAGE analysis of the His-tagged Jermuk-LamM. Line 1 corresponds to the cell lysate of *E. coli* C43(DE3) after induction; line 2 crude extract; line 3 flow through, line 4 eluate after Ni-NTA affinity chromatography; line 5 purified enzyme after size exclusion chromatography; line 6 molecular weight marker.

### Substrate specificity of Jermuk-LamM

The substrate specificity of Jermuk-LamM was tested towards a number of complex carbohydrates at 65 °C (**Table 1**). The highest activity of the enzyme was measured towards laminarin with 217 ± 8.1 U/mg. Compared to that, its highest activity towards amorphous curdlan was 74 ± 17.39 U/mg. For undissolved curdlan, the highest activity was 35 ± 21.59 U/mg. The reason for the relatively low activity of the enzyme against curdlan is unclear, but it may be due to poor dispersal of the substrate in the assay tubes. In comparison to β-1,3-glucans, the specific activities of Jermuk-LamM towards mixed-linked glucans such as barley β-glucan and lichenin were 128 ± 8.6 U/mg and 27 ± 3.55 U/mg, respectively. The specific activity towards lichenin was lower than towards the other tested substrates. Hydrolysis of the β-1,4-glucan substrates (CMC, xyloglucan and xylan) was not observed, further supporting the hypothesis that this enzyme acts primarily as β-1,3-glucanase.

**Table 1.** Substrate specificity of Jermuk-LamM.

| Substrate | Specific activity (U/mg) |
| --- | --- |
| Laminarin | $217.7 \pm 8.1$ |
| Curdlan * | $74.2 \pm 17.4$ |
| Curdlan | $35.5 \pm 21.6$ |
| Barley β-glucan | $128.3 \pm 8.6$ |
| Lichenin | $27.2 \pm 3.6$ |
| Xyloglucane | n.d** |
| Xylane | n.d |
| CMC + | n.d |

\* Amorphous curdlan, diluted in NaOH and neutralized
\*\* Not detectable

### Degradation pattern and enzyme kinetics

The hydrolysis products of laminarin and barley β-glucan were investigated by HPLC analysis. Hydrolysis of laminarin (Figure 4a) mainly resulted in glucose (DP1) and laminaribiose, glucose-mannitol units (DP2), and to a lesser extent laminaritriose (DP3) and higher oligosaccharides. The degradation pattern of barley β-glucan was different (Figure 4b). Laminaritriose was the major product and to a lesser extent laminaritetrose (DP4) and other oligosaccharides were detected. The produced oligosaccharides indicated that Jermuk-LamM is a β-1,3-glucanase with endo-acting mode. Moreover, the shifted product variation with barley β-glucan suggests that the enzyme hydrolyzes the β-1,3-glycosidic linkages in mixed-linked glucans and to a lesser extent the β-1,4-building blocks. It is very important to note that after 18 hours reaction both substrates (0.25 gram each) were completely hydrolyzed with 1 U of enzyme․

**Figure 4.**
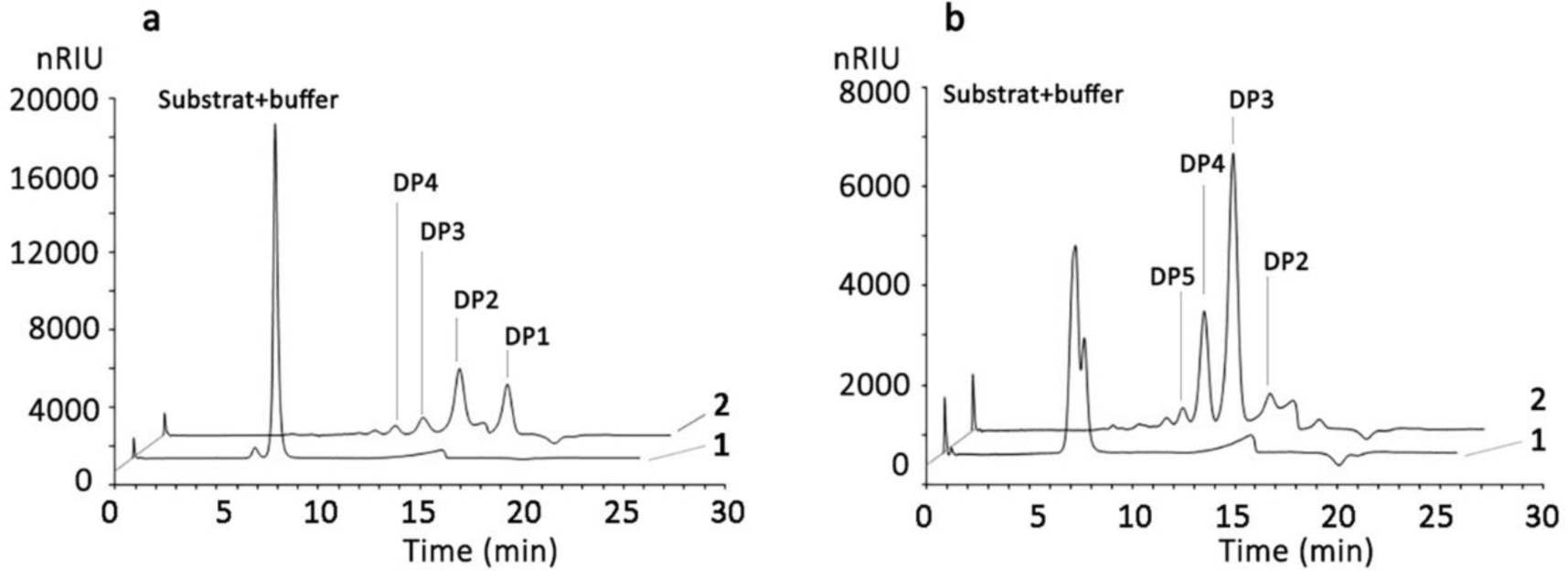
HPLC analysis of products observed from polysaccharide degradation. Hydrolysis products from laminarin (a) and barley β-glucan (b) after 18 h of incubation at 55 °C. DP: degree of polymerization. DP1: glucose, DP2 laminaribiose, DP3 laminaritriose, DP4 laminaritetrose, DP5 laminaripentose. 1: substrate; 2: substrate + Jermuk-LamM.

### Effects of pH and Temperature

The effect of pH was analyzed in a range of pH 2 to 9 (Figure 5). Jermuk-LamM showed activity in a range between pH 4.5 and 7.5. Maximum activity was detected around pH 6.

**Figure 5.**
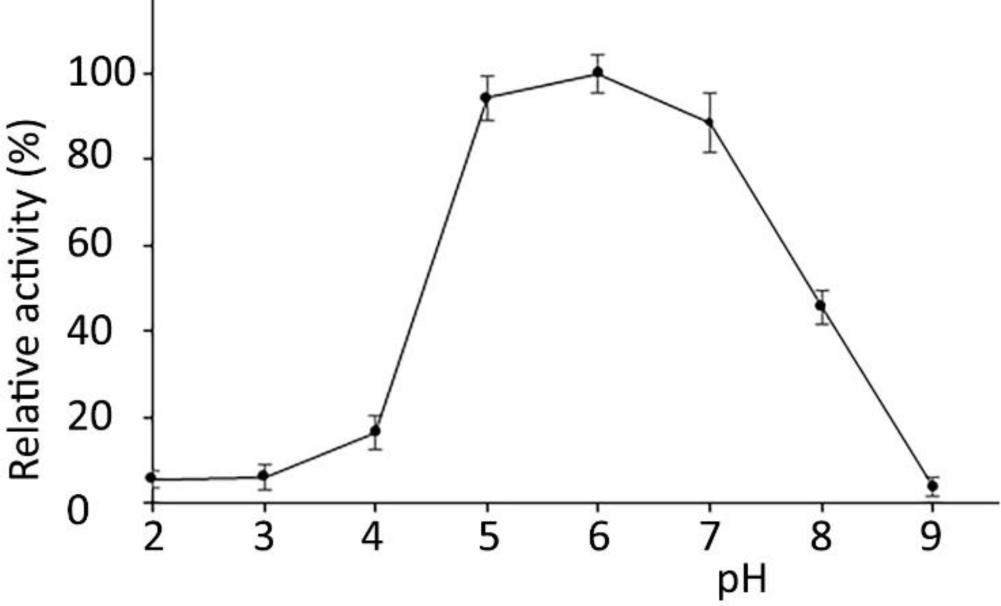
Influence of pH on the activity of recombinant Jermuk-LamM towards laminarin Connections between dots are solely for better visualization of the enzyme’s activity range.

To examine the temperature optimum, Jermuk-LamM activity was measured in a temperature range from 30 °C to 80 °C. Maximum activity of the enzyme was measured at 65 °C (Figure 6a). To examine the temperature sensitivity of the enzyme, residual activities were measured after 10 min of incubation at different temperatures ranging from 30 °C to 80 °C (Figure 6b). The enzyme lost 50 % of its initial activity after incubation at 65 °C for 10 min. To analyse the temperature stability at selected temperatures, the enzyme was pre-incubated at 55 °C, 60 °C and 65 °C for 24 hours (Figure 6c). Residual activity was obtained in relation to initial activity prior to pre-incubation. The enzyme exhibited more than 50 % residual activity over 24 h at 55 °C. At 60 °C, the enzyme had a half-life time of 3 h. After incubating the enzyme at 65 °C for 1 hour, less than 10 % residual activity was detected.

**Figure 6.**
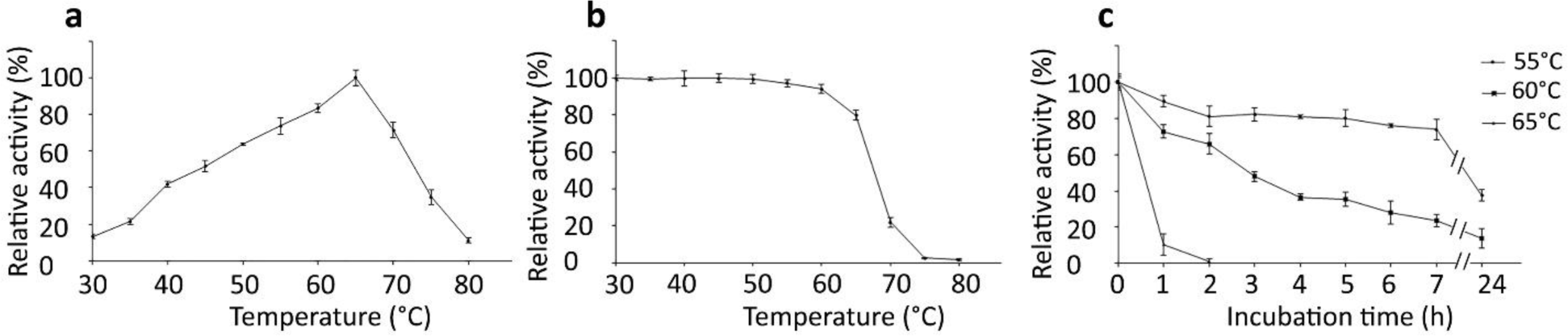
Temperature profile of Jermuk-LamM. (a) Temperature spectrum in a range of 30-80 °C; (b) thermal stability after 10 minutes incubation at a temperature in the range of 30-80 °C, (c) influence of temperature (55 °C circle, 60 °C square, 65 °C triangle) on the stability of Jermuk-LamM. Connections between dots are solely for better visualization of the enzyme’s activity range.

The effect of various additives on the activity of Jermuk-LamM was investigated. Among the tested metal ions (5 mM), only Fe^2+^ and Zn^2+^ led to significant inhibition of 86 % and 60 %, respectively. Fe^3+^, Al^3+^, and Cu^2+^ reduced the activity of the enzyme to 30 %. In contrast, Mn^2+^ had a positive effect on enzyme activity (Figure 7a). Inhibition was observed by the addition of 5 mM SDS, CHAPS and sodium-periodate (Figure 7b). The reducing agents dithiothreitol (DTT) and β-mercaptoethanol showed only slight inhibitory effects on the enzyme’s activity at 5 mM concentration.

**Figure 7.**
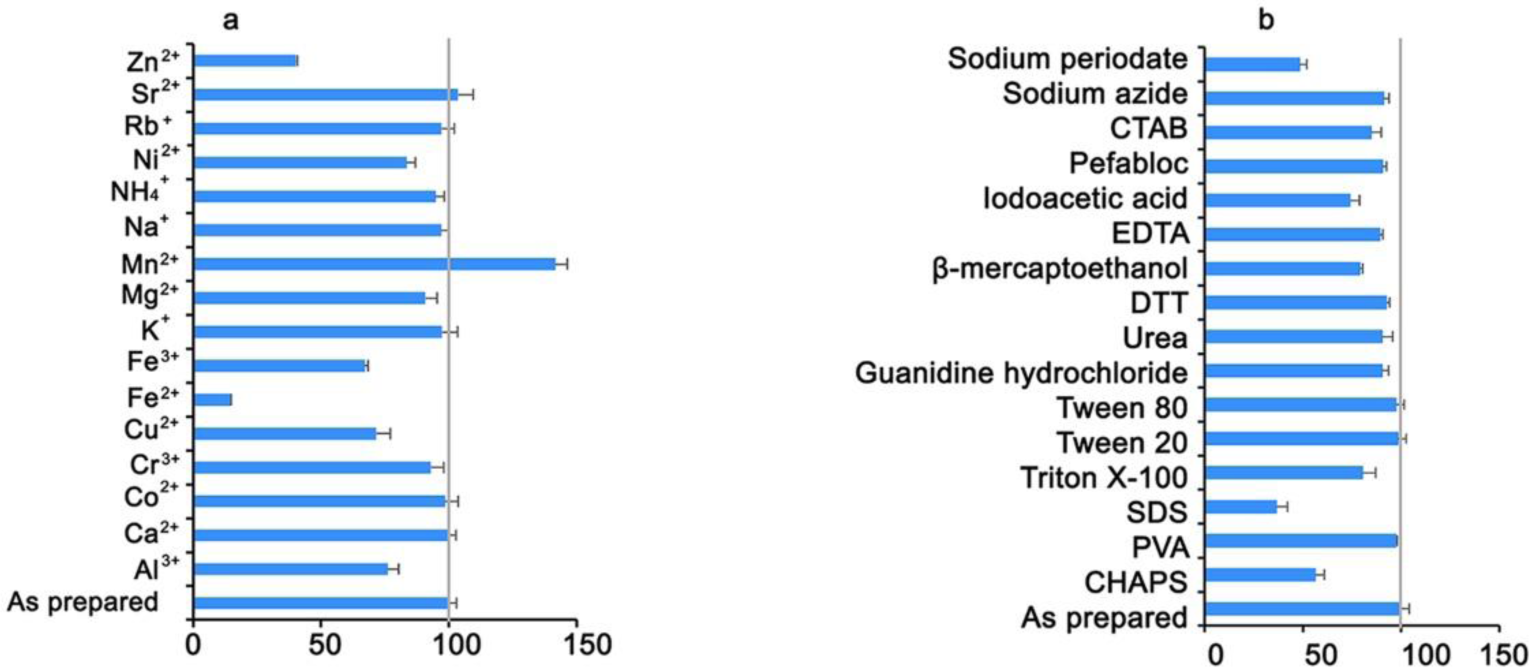
Effect of cations and chemical compounds on the activity of Jermuk-LamM. A) The activity of the recombinant Jermuk-LamM enzyme was assessed after one-hour of incubation at room temperature in the presence of 5 mM of different metals (a) and additives (b). A blank experiment was conducted for each metal and additive. A specific activity of 217 U/mg obtained without additives was defined as 100 % activity.

## Discussion

In the light of current environmental challenges, such as climate change and the depletion of fossil fuel reservoirs, a global transition to biodegradable and renewable products is highly demanded. Recently, the degradation of macroalgae biomass, as a renewable source of energy, has become attractive, and intensive efforts are underway to develop valorization technologies [78, 79]. Consequently, high-value bioactive compounds can be extracted from macroalgae biomass, finding utility as feed and food additives [80, 81]. Additionally, a large variety of bioactive compounds originating from algae finds application in both the prevention and treatment of various diseases, as well as in the pharmaceutical and cosmetics industries [82, 83]. Furthermore, simple monomers can be derived from the polysaccharides and proteins within macroalgal biomass, serving as a valuable source of fermentable nutrients, etc. [84]. In order to make macroalgae biomass applicable for these purposes, effective and sustainable processing methods need to be developed in which the role of enzymes is vital [85].

A novel β-1,3-glucanase was identified from Jermuk hot spring metagenome of Armenia using sequence-based metagenomics approach was used to identify putative 1,3-β-glucanase sequences from the Jermuk metagenome.

NCBI Blast search results indicated that the ORF Jermuk-lamM has a marine microbial origin showing the highest identity to <u>Candidatus</u> *Marinimicrobia* bacterium. This strain is an uncultivated bacterium and widespread in sub-euphotic areas and particularly abundant in minimum oxygen zones. The information on *Marinimicrobia* and their metabolism is sparse, making the biogeochemical influence of this group challenging to predict. Available genome information revealed a tremendous diversity of *Marinimicrobia* in marine environments where they play an important role in cycling of sulfur and nitrogen [86]. Based on our knowledge this is the first β-1,3-glucanase characterised among the *Marinimicrobia* phylum and could provide useful initial information towards revealing their metabolic functions. The detection of this strain in the Jermuk hot spring probably is attributed to the spring’s relatively high concentration of dissolved minerals [87]․

This enzyme has the highest identity of only 48 % to characterised laminarinases. Such a low similarity percentage to characterised enzymes suggests that Jermuk-LamM is quite different and unique from the laminarinases characterised so far.

β-1,3-linkage hydrolyzing enzymes generally show preferences either for β-1,3-glucans (laminarinase EC 3.2.1.39) or mixed-linkage glucans (β −1,3(4)-glucanase (EC 3.2.1.6). Even though Jermuk-LamM clusters with endo-1,3(4)-beta-glucanase of *Acetivibrio thermocellus* (Figure 1) they are completely different in their substrate specificity. Mainly endo-1,3(4)-beta-glucanase of *Acetivibrio thermocellus* shows more than 7-times higher specific activity to mixed linkage glucans than to laminarin [75]. Meanwhile like other bacterial endo-β-1,3-glucanases [36, 88, 48] except LamR from *Rhodothermus marinus* [21], and BglF from *Nocardiopsis* sp. strain F96 [39] Jermuk-LamM degrades laminarin (100 %) and to a lesser degree lichenan (10 %) (**Table 1**). The substrate specificity profile of Jermuk-LamM closely resembles that of FLamA and FLamB identified in thermophilic *Fervidobacterium* sp. These enzymes share a common order of substrate preferences, prioritizing Laminarine > Curdlan > Barley β-glucan > Lichenin. Remarkably, Jermuk-LamM displays less than 43 % similarity [89]. The enzyme cleaved the substrates randomly, yielding glucose, laminaribiose, and laminaritriose as end products (Figure 4). Although the substrate specificity profiles are quite similar, there are notable distinctions in the end products resulting from the hydrolytic reactions catalysed by Jermuk-LamM and the laminarinases of Fervidobacterium sp. In particular, as a result of barley β-glucan hydrolises, Jermuk-LamM produces minimal glucose, while FLamA and FLamB generate glucose as one of the primary end products [89]. Thus, based on the biochemical characteristics, substrate specificity and product pattern we classify Jermuk-LamM as a typical endo-β-1,3-glucanase of the GH16_3 subfamily (EC 3.2.1.39) [90] together with TmβG from the bacterium *Thermotoga maritima* MSB8 [43] and LamA from the archaeon *Pyrococcus furiosus*, respectively [91].

The molecular weight of laminarinases is quite diverse and mainly depends on the composition of different domains. The molecular weight of Jermuk-LamM (30 kDa) is close to those of ULam111 (27 kDa) from *Flavobacterium* sp. [46], and FLamA and FLamB (34.9 kDa and 34.1 kDa, respectively) from *Fervidobacterium* sp [89]; β-glucanase (27 kDa) of *Bacillus halodurans* C-125 [38] and archaeal LamA (30 kDa) from *Pyrococcus furiosus* [91]. It was reported that substrate preferences were dependent on the carbohydrate-binding domain or an extra loop/short amino acid sequence that are well-suited for binding the β-1,3-glucan [49, 77, 76, 47, 45]. Surprisingly, Jermuk-LamM does not contain any carbohydrate-binding domain or similar peptide but has noticeable substrate preference to laminarin (**Table 1**). ULam111 also lacks the flexible loop at the entrance of substrate recognition, however, the disulfide bond of two cysteine amino acids regulates it [46]. With respect to laminarin as a preferred substrate Jermuk-LamM was found to be similar to LamA [36], FLamA and FLamB [89], ULam111 [46] and *Zg*LamA [49].

Regarding the temperature maxima of these enzymes, with 65 °C Jermuk-LamM has a temperature maximum close to other examined β-1,3-glucanases, such as β-glucanase (60 °C) of *Bacillus halodurans* C-125 [38], BglF (70 °C) of *Nocardiopsis* sp. strain F96 [39] and laminarinase (65 °C) of *Clostridium thermocellum* [92]. Concerning the pH profile, the enzyme retains at least 80 % of its activity in a pH range from 5 to 7 with maximum activity at pH 6. Similar results have been obtained for 1,3-β-glucanase (pH 5.5) from *Streptomyces sioyaensis* [35], LamR (pH 5.5) from *Rhodothermus marinus* ITI278 [21], LamA (pH 6) from *Pyrococcus furiosus* [36], and laminarinase (pH 6.5) from *Clostridium thermocellum* [92].

Based on the experimental findings, Jermuk-LamM is a novel enzyme with significant potential for industrial applications. Due to its substrate specificity and hydrolytic activity to produce fermentable sugars Jermuk-LamM can be successfully applied for macroalgae biomass processing such as saccharification and degradation of storage polysaccharides. An essential advantage of this enzyme lies in its resilience to metal ions, detergents, and other chemicals. This characteristic significantly broadens the potential applications of this enzyme across diverse industrial conditions. The characteristics of Jermuk-LamM give a possibility for an application not only in different industries but also make this enzyme a promising candidate for enzymatic cocktail formulation. This allows the development of sustainable technologies as an alternative to acid and heat treatment of macroalgal biomass.

## Acknowledgements

The authors would like to thank Dr. Philip Busch and Henning Piascheck for sampling, and Dr. Christian Schäfers for setting up the bioinformatics pipeline.

## Author Contributions

Study conceptualization: AP, AK, GA

Investigation: Cloning and characterisation of enzyme: AP; HPLC analyses: ChB

Writing – Original Draft: AP

Writing – Review and Editing: AP, AK, GA, ChB

## Data availability Statement

All data used to prepare this manuscript are available as supplementary materials or deposited at publicly accessible databases.

