## Supplemental for "Molecular Cloning and Biochemical Characterisation of a Novel Acidic Laminarinase Derived from Jermuk Hot Spring Metagenome"

**Contents**

**Supplementary Figures**

**Supplementary Figure 1.** The analysis of Jermuk-LamM protein domain architecture.

**Supplementary Figure 2.** Structure of Jermuk-LamM

**Supplementary Tables**

**Supplementary Table 1.** Purification table for recombinant Jermuk-LamM from *E. coli* C43(DE3).

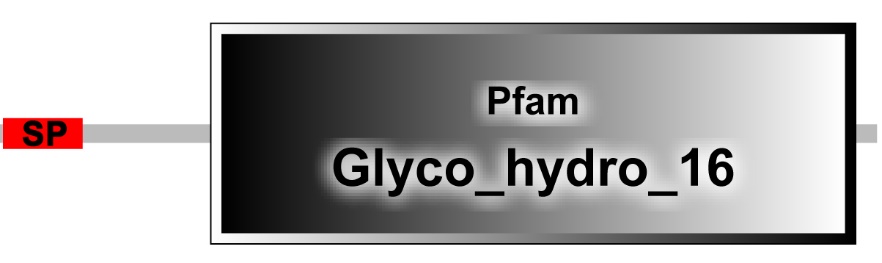

**Supplementary Figure 1. The analysis of Jermuk-LamM protein domain architecture.**

The SMART (Simple Modular Architecture Research Tool <https://smart.embl.de>) has predicted the presence of a signal peptide (SP) containing a 27-amino acid sequence and a glycosyl hydrolase family 16 catalytic domain (CH16), containing 211 amino acids (starting from Gly69). These two domains are linked by a 41-amino acid peptide.

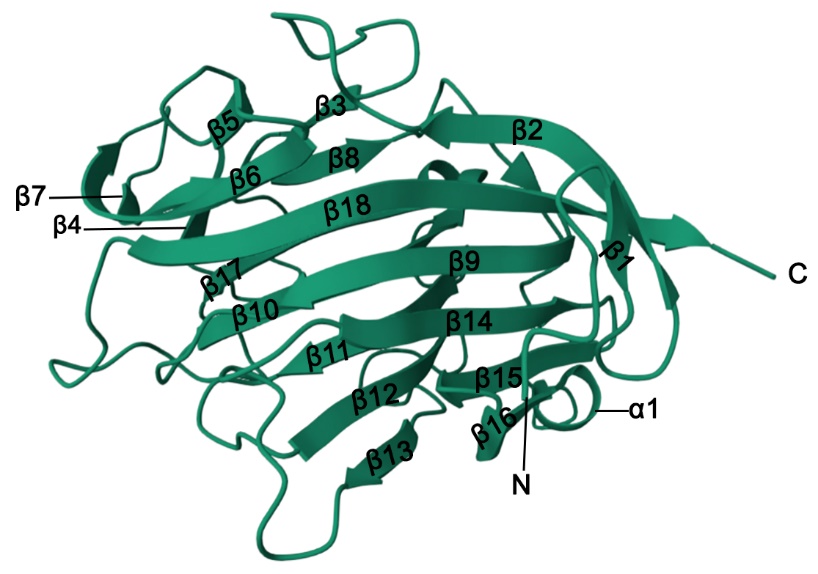

**Supplementary figure 2.** The 3D structure modeling of Jermuk-LamM. The structure was modeled using endo-beta-1,3-glucanase (BglF) of alkaliphilic *Nocardiopsis sp.* strain F96 (PDB: 2hyk) as a template showing 52 % identity (98% of residues modelled at >90% confidence). 18 β-strands and an α-helice are labelled.

**Supplementary tables**

**Supplementary Table 1. Purification table for recombinant Jermuk-LamM from *E. coli* C43(DE3).**

| Purification steps | Total protein (mg) | Total activity (U) | Specific activity (U/mg) | Yield (%) |
| --- | --- | --- | --- | --- |
| Crude enzyme | 1468.5 | 210.3 | 0.14 | 100 |
| His_Tag | 2.34 | 43.7 | 18.67 | 20.7 |
| Sephacryl S-100 | 0.13 | 28.6 | 228.73 | 13.6 |
